# A new coactivator complex required for retinoic acid-dependent regulation of embryonic symmetry

**DOI:** 10.1101/089201

**Authors:** Goncalo C. Vilhais-Neto, Marjorie Fournier, Jean-Luc Plassat, Mihaela E. Sardiu, Anita Saraf, Jean-Marie Garnier, Mitsuji Maruhashi, Laurence Florens, Michael P. Washburn, Olivier Pourquié

## Abstract

Bilateral symmetry is a striking feature of the vertebrate body plan organization. Vertebral precursors, called somites, provide one of the best illustrations of embryonic symmetry. Maintenance of somitogenesis symmetry requires Retinoic acid (RA) and its coactivator Rere/Atrophin2. Here, using a proteomic approach we identify a protein complex, containing <u>W</u>dr5, <u>H</u>dac1, <u>H</u>dac2 and R<u>ere</u> (named WHHERE), which regulates RA signalling and controls embryonic symmetry. We demonstrate that Wdr5, Hdac1 and Hdac2 are required for RA signalling *in vitro* and *in vivo*. Mouse mutants for *Wdr5* and *Hdac1* exhibit asymmetrical somite formation characteristic of RA-deficiency. We also identify the Rere-binding histone methyltransferase Ehmt2/G9a, as a RA coactivator controlling somite symmetry. Upon RA treatment, WHHERE and Ehmt2 become enriched at RA target genes to promote RNA Polymerase II recruitment. Our work identifies a novel protein complex linking key epigenetic regulators acting in the molecular control of embryonic bilateral symmetry.

---

The development of bilaterally symmetrical structures such as limbs or somites takes place concomitantly with the asymmetric formation of internal organs such as heart, gut and liver. Whereas the pathway responsible for establishing left-right identity in the embryo begins to be well understood^1^, little is known about the mechanisms controlling embryonic symmetry. Retinoic acid (RA) is a derivative of vitamin A, signalling via a heterodimeric RAR/RXR nuclear receptor transcription factor^2-4^. In the absence of RA, the heterodimer binds target genes together with the SMRT and NCoR corepressor complexes and histone deacetylases such as Hdac3 to silence gene expression. When the RA ligand binds to RAR, the corepressors are replaced by a set of coactivators including histone acetyltransferases, contributing to active transcription of RA target genes^5, 6^. In the absence of RA signalling in the mouse embryo, somite formation becomes asymmetrical, showing a significant delay on the right side^7^. A similar somite desynchronization phenotype is also observed in mutants for the protein Rere (or Atrophin2) which acts as a coactivator for RA signalling^8^.

In order to understand the mechanism of action of Rere in the RA pathway controlling somite symmetry, we first set out to identify Rere-interacting proteins in the mouse mesoderm. To that end, we generated a transgenic mouse line, which allows the conditional expression of a tagged version of Rere containing two HA epitopes at the C-terminal end of the protein (Rere-HA). A *Rere-HA* construct preceded by a *LoxP-STOP-LoxP* cassette was introduced into the *Rosa26* locus by homologous recombination in mouse embryonic stem (ES) cells. We then used these cells to generate a <u>*R*</u>*osa26-LoxP*-<u>*S*</u>*TOP-LoxP-Rere-HA* mouse line (*RS-Rere-HA* line). Whereas *Rere* mutants (*Rere*^*om/om*^) die around E9.5 with defects in forebrain, heart and a right-side specific delay in somite formation^8, 9^, expression of at least one *Rere-HA* allele in the mutant *Rere*^*om/om*^ background led to morphologically normal embryos (Supplementary Fig. 1a-c). Therefore, the tagged Rere-HA protein is functional in vivo. To direct expression of Rere-HA to the mesoderm, *RS-Rere-HA* mice were crossed to the *T-Cre* mouse line^10^. We prepared whole cell protein extracts from approximately 600 *RS-Rere-HA;T-Cre* embryos, and performed affinity purification using anti-HA antibodies under high or low salt conditions (Fig. 1a). To identify the immunoprecipitated proteins, eluted fractions were submitted to mass spectrometry analysis using the multidimensional protein identification technology (MudPIT)^11^. A set of 105 common proteins was found between the different immunopurification conditions (Supplementary Fig. 1d and Supplementary Table 1). Hierarchical clustering analysis of the 105 proteins based on the normalized spectral abundance factor (NSAF)^12^ in the different immunoprecipitation conditions identified three abundant proteins tightly clustering with Rere (Supplementary Fig. 1e,f). These include Rere’s known binding partners Hdac1 and Hdac2 as well as a novel partner, Wdr5^13-16^ (Fig. 1b and Supplementary Fig. 2a). To estimate relative protein levels, we compared the NSAF values for each protein^12^. NSAF for Rere, Hdac1 and Hdac2 were similar while it was three times higher for Wdr5 (Fig. 1b). These results suggest that the proteins Rere, Hdac1, Hdac2 and Wdr5 can interact in the mesoderm.

**Figure 1.**
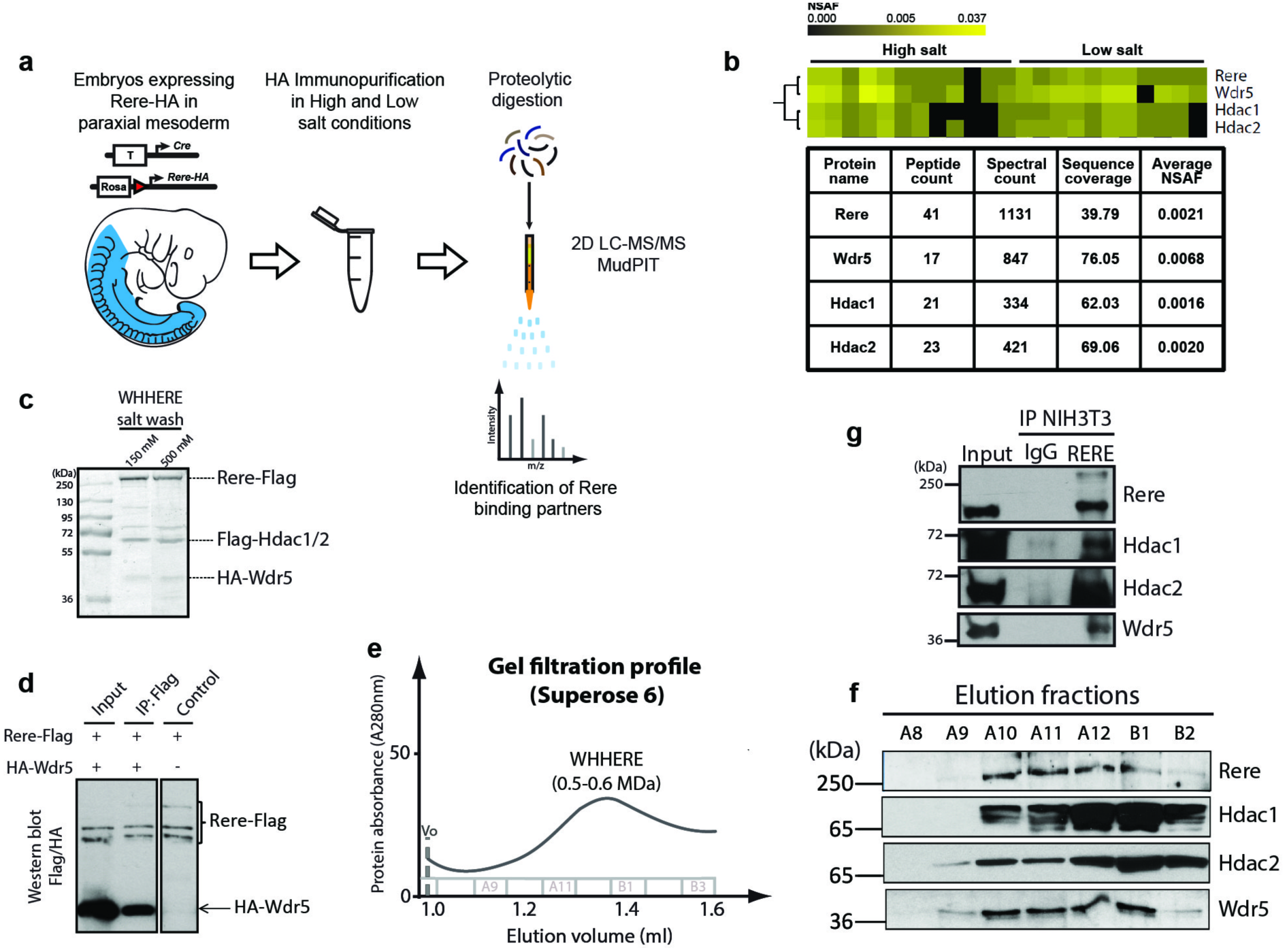
Proteomic identification of the WHHERE protein complex containing <u>W</u>dr5, <u>H</u>dac1, <u>H</u>dac2 and R<u>ere</u>. (**a**) Schematic workflow for the proteomic strategy used to identify Rere-associated proteins. (**b**) Hierarchical clustering of the NSAF values showing Wdr5, Hdac1 and Hdac2 clustering with Rere identified by MudPIT. Columns correspond to different immunoprecipitations conditions. Table showing the peptide count, spectral count, sequence coverage (%) and average NSAF values for Rere, Wdr5, Hdac1 and Hdac2. (**c**) SDS-PAGE gel stained with Coomassie blue after Flag immunopurification at low (150 mM KCl) and high (500 mM KCl) salt wash of the recombinant WHHERE complex from baculoviruses infected cells co-expressing Rere-Flag, Flag-Hdac1, Flag-Hdac2 and HA-Wdr5. Identification of the different components was confirmed by LC-MS/MS analysis of the corresponding gel bands. (**d**) Flag immunoprecipitation from extracts of cells infected with baculoviruses expressing HA-Wdr5 and Rere-Flag. Flag and HA western blot. The three bands recognized by the anti-Flag antibodies correspond to Rere and its degradation products. (**e**) Gel filtration chromatography profile of the WHHERE complex purified from baculoviruses infected cells co-expressing Rere-Flag, Flag-Hdac1, Flag-Hdac2 and HA-Wdr5. (**f**) Immunoblot analysis for Rere, Hdac1, Hdac2 and Wdr5 showing the gel filtration chromatography elution fractions from A8 to B2. (**g**) Rere immunoprecipitation from NIH3T3 cells identifying the endogenous WHHERE complex components. Rere, Hdac1, Hdac2 and Wdr5 were detected after western blot analysis of Rere immunoprecipitation eluates and compared to a negative control immunoprecipitation (IgG) and input extract.

To validate the identification of Wdr5 as a novel interacting partner of Rere, Hdac1 and Hdac2, we co-expressed tagged versions of the four proteins (Rere-Flag, Flag-Hdac1, Flag-Hdac2 and HA-Wdr5) using a baculovirus-insect cell expression system. After Flag immunoprecipitation of Rere, Hdac1 and Hdac2, HA-Wdr5 was detected in the eluates, as confirmed by LC-MS/MS analysis of the Coomassie stained gel bands (Fig. 1c and Supplementary Fig. 2b). The four proteins still co-purified together at high salt concentration (500 mM KCl) suggesting the existence of a stable protein interaction network between Rere, Hdac1, Hdac2 and Wdr5 (Fig. 1c). By co-immunoprecipitation of baculovirus-expressed Rere-Flag and HA-Wdr5, we could demonstrate that Rere binds directly to Wdr5 (Fig. 1d). However Wdr5 does not bind directly to Hdac1 or Hdac2 in high or low salt wash (Supplementary Fig. 2c), suggesting that Rere acts as a scaffolding component binding Hdac1/Hdac2 and Wdr5. To analyse whether the four co-immunopurified proteins form a stable protein complex, we carried out gel filtration chromatography followed by western blot and mass spectrometry analysis. All four proteins co-eluted together in a fraction corresponding to a high molecular weight complex of 0.5-0.6 MDa (Fig. 1e,f and Supplementary Fig. 2d,e). This molecular weight is consistent with the abundance predicted by the NSAF values of the proteomic analysis. Additionally, Hdac1, Hdac2 and Wdr5 co-immunoprecipitated with the endogenous Rere in NIH3T3 cells further supporting the existence of such a protein complex (Fig. 1g). Altogether, these results suggest that <u>W</u>dr5, <u>H</u>dac1, <u>H</u>dac2 and R<u>ere</u> can form a stable protein complex, which we called the WHHERE complex.

To test the coactivator properties of WHHERE on RA signalling, we transfected NIH3T3 fibroblast cells with expression vectors coding for either *Rere, Wdr5, Hdac1* or *Hdac2*, together with a RA reporter containing the well-characterized Retinoic Acid Response Element (RARE) of the known RA target *Retinoic Acid Receptor beta* (*Rarβ)*^17^, driving luciferase expression. In the presence of RA, overexpression of each of the four WHHERE complex proteins increased RA reporter expression (Fig. 2a). Furthermore, co-transfection of *Hdac1* and *Hdac2* cooperates to activate RA signalling (Fig. 2b). Conversely, treatment of fibroblast cultures with siRNAs against *Rere, Hdac1* or *Wdr5* in the presence of RA decreased RARE-Luciferase reporter activity (Fig. 2c and Supplementary Fig. 3a-c). Treatment with *Hdac2* siRNA led to an increase of RARE-Luciferase reporter activity (Fig. 2c and Supplementary Fig. 3d), which might be explained by a stabilization of Hdac1 (due to the decrease of Hdac2), potentially resulting in an increase in RA signalling^18, 19^. Consistent with this possibility, the double knockdown of *Hdac1* and *Hdac2* further decreased RA signalling compared to *Hdac1* siRNA alone (Fig. 2d). Furthermore, *Hdac1* or *Hdac1/Hdac2* depletion reduced RA activation mediated by *Rere* and *Wdr5* (Fig. 2e,f). Inhibition of deacetylase enzymatic activity with a range of chemical inhibitors decreased RA signalling (Fig. 2g)^20^. In line with this, overexpression of *Rere* or *Wdr5* did not lead to a significant increase in RA signalling when cells were treated with the HDAC inhibitors Trichostatin A (TSA) or Sodium Butyrate (SB) (Fig. 2h,i). Hdac1 and Hdac2 have been shown to bind Rere N-terminal region^9, 13, 14^. In the presence of RA, overexpression of the N-terminal region of Rere (N-Rere) strongly increased RA signalling whereas no activation could be seen with Rere C-terminal region (Rere-C) (Fig. 2j). The activation by N-Rere was dependent on deacetylase activity since TSA or SB treatment strongly decreased N-Rere dependent RA signalling (Fig. 2k). Overall, these results show that the WHHERE complex proteins Rere, Wdr5, Hdac1 and Hdac2 act to trigger the RA pathway activation in NIH3T3 cells. Moreover, activation of RA signalling by the WHHERE complex depends on Hdac1 and Hdac2 deacetylase activity.

**Figure 2.**
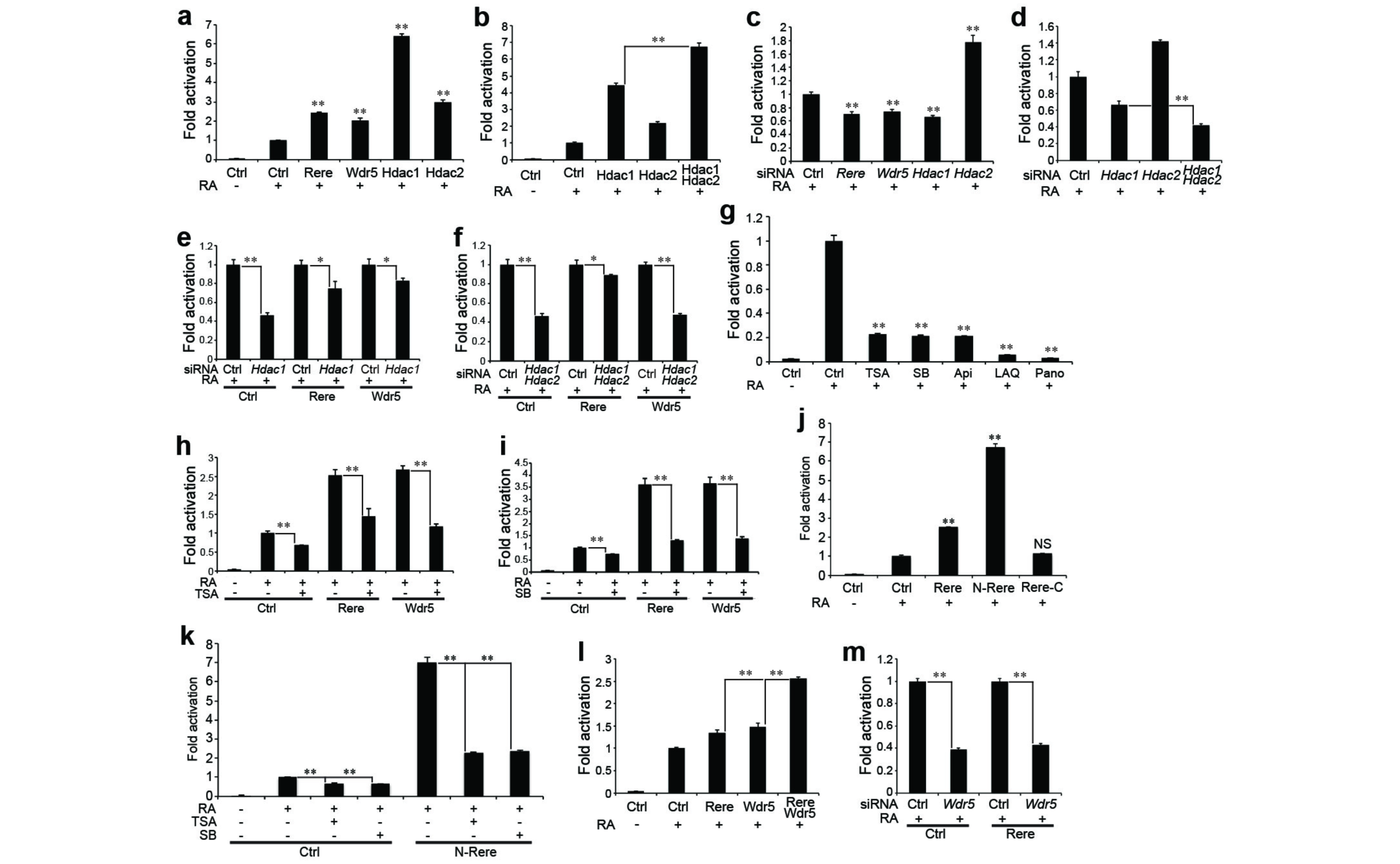
The WHHERE complex acts as a coactivator for retinoic acid signalling. (**a-k**) RARE-Luciferase activity from NIH3T3 cells treated or not with 1 μM RA for 20 hours. (**a**) Cells transfected with expression plasmids containing *Rere, Wdr5*, *Hdac1* or *Hdac2* (n = 4). (**b**) Cells transfected with expression plasmids containing *Hdac1*, *Hdac2* or both (n = 4). (**c**) Cells treated either with siRNA for *Rere, Wdr5*, *Hdac1* or *Hdac2* (n = 4). (**d**) Cells treated with siRNA for *Hdac1*, *Hdac2* or both (n = 4). (**e**) Cells overexpressing *Rere* or *Wdr5* and treated with siRNA against *Hdac1* (n = 4). (**f**) Cells overexpressing *Rere* or *Wdr5* and treated with siRNA against both *Hdac1* and *Hdac2* (n = 4). (**g**) Cells treated with the HDAC inhibitors Trichostatin A (TSA) (60 nM), Sodium butyrate (SB) (3 mM), Apicidin (Api) (300 nM), LAQ824 (LAQ) (60 nM) and Panobinostat (Pano) (30 nM) (n = 4). (**h**) Cells overexpressing *Rere* or *Wdr5* and treated with TSA (30 nM) (n = 4). (**i**) Cells overexpressing *Rere* or *Wdr5* and treated with SB (1.5 mM) (n = 4). (**j**) Cells transfected with expression plasmids containing *Rere*, *N-Rere* (*Rere* N-terminal domain) or *Rere-C* (*Rere* C-terminal domain) (n = 3). (**k**) Cells overexpressing *N-Rere* (*Rere* N-terminal domain) and treated with TSA (30 nM) or SB (1.5 mM) (n = 4). (**l**) Cells transfected with expression plasmids containing *Rere*, *Wdr5* or both (n = 4). (**m**) Cells overexpressing *Rere* and treated with siRNA for *Wdr5* (n = 4). In all graphs data represent mean ± s.e.m‥ NS - not significant, \**P* < 0.05 and \*\**P* < 0.01.

To analyze WHHERE-dependent RA regulation *in vivo*, we characterized the phenotype of mouse embryos mutant for the different components of the complex. *Hdac1* mutants exhibit a variety of developmental defects similar to *Rere*^*om/om*^ embryos^21^. We introduced the *RARE-LacZ* reporter^22^ in *Hdac1*-null embryos and observed a strong downregulation of LacZ expression, similar to that seen in *Rere* mutants (Fig. 3a-c). *Hdac2* mutants do not show developmental defects and can survive until the perinatal period^21^. To analyze Wdr5 function *in vivo* we first set out to generate a conditional knock-out mouse line by introducing *LoxP* sites flanking exons 2 to exon 4 (Supplementary Fig. 4a). No heterozygous embryos were recovered at E8.5 after crossing with an ubiquitous Cre suggesting that the *Wdr5* mutation is heterozygous lethal at early stages. To circumvent this issue, we crossed mice homozygous for the *Wdr5* conditional allele (*Wdr5*^*fl/fl*^) to the mesoderm-specific *T-Cre* line^10^. Removal of a single *Wdr5* allele in *Wdr5*^*fl/*+^*;T-Cre;RARE-LacZ* embryos was sufficient to strongly decrease beta-galactosidase staining in the mesoderm (Fig. 3d). In contrast, no significant downregulation of Notch target genes such as *Lfng* or *Hes7* was detected in these mutants suggesting the Notch-dependent somite segmentation clock appears normal (Supplementary Fig. 4b-e). Altogether, these results support the function of the WHHERE complex in the control of RA signalling in the embryo.

**Figure 3.**
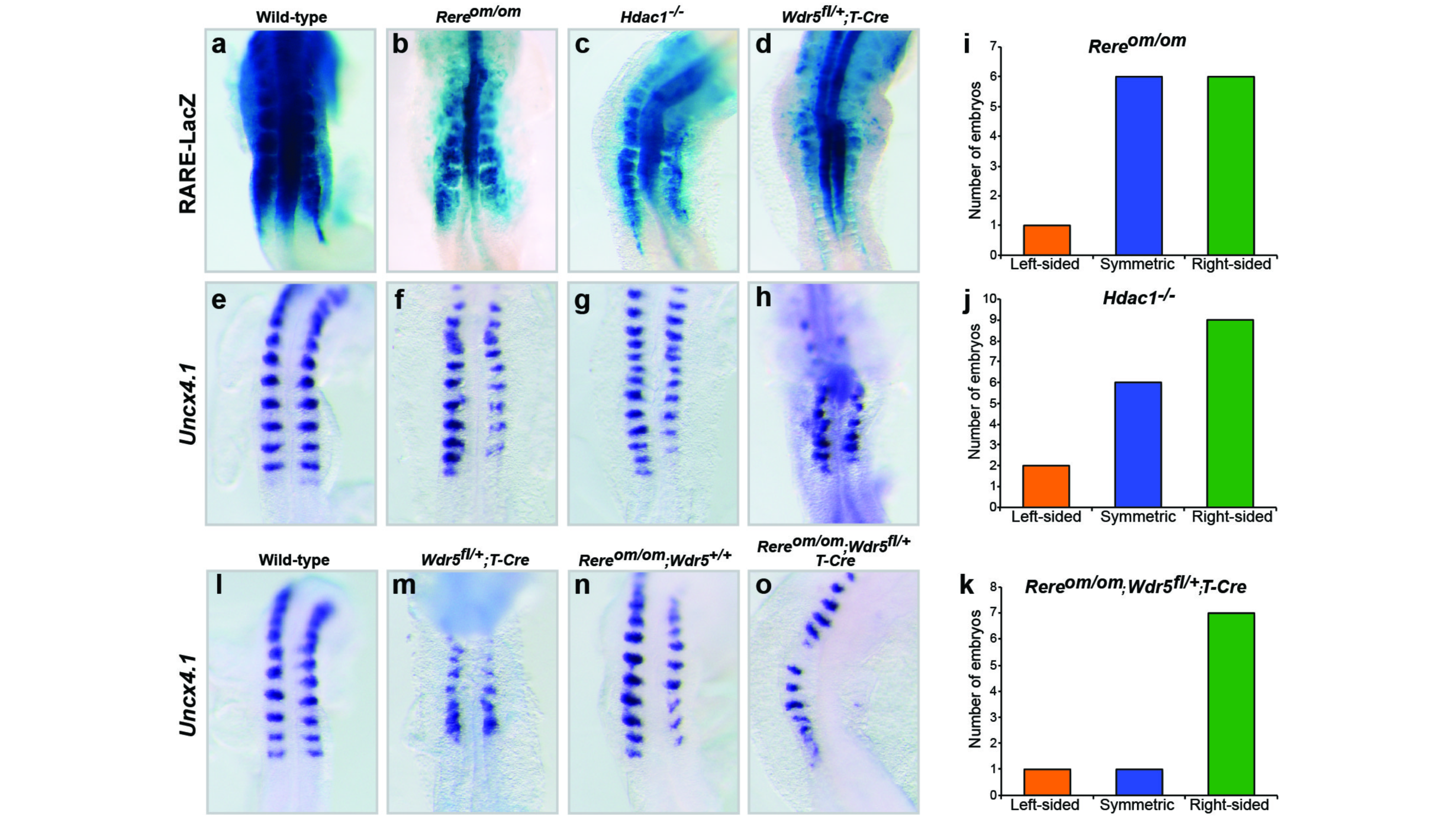
The WHHERE complex controls somite bilateral symmetry. (**a-d**) RARE-LacZ expression in wild-type (**a**), *Rere*^*om/om*^ (**b**), *Hdac1*^−*/*-^ (**c**) and *Wdr5*^*fl/*+^*;T-Cre* (**d**) embryos at E8.75-E9.0 (dorsal views). (**e-h**) *In situ* hybridization showing somites labeled with *Uncx4.1* in wild-type (**e**), *Rere*^*om/om*^ (**f**), *Hdac1*^-*/*-^ (**g**) and *Wdr5*^*fl/*+^*;T-Cre* (**h**) embryos at E8.75-E9.0 (dorsal views). (**i-k**) Graphs representing the number of 7- to 15-somite stage embryos with left-sided (orange), symmetric (blue) or right-sided (green) delay in somite formation: *Rere*^*om/om*^ (**i**), *Hdac1*^-*/*-^ (**j**) and *Rere*^*om/om*^;Wdr5^*fl/*+^*;T-Cre* (**k**). (**l-o**) *In situ* hybridization for *Uncx4.1* in wild-type (**l**), *Wdr5*^*fl/*+^*;T-Cre* (**m**), *Rere*^*om/om*^;*Wdr5*^+*/*+^ (**n**) and *Rere*^*om/om*^;*Wdr5*^*fl/*+^*;T-Cre* (**o**) embryos at E8.75-E9.0 (dorsal views).

We next examined the effect of null mutations of members of the WHHERE complex on somite symmetry. In nearly half of the *Hdac1*^-*/*-^ embryos, we observed a right side delay of somite formation resembling the defect observed in *Rere*^*om/om*^ and *Raldh2*^-*/*-^ mutants^7, 8^ (Fig. 3e-g,i,j and Supplementary Fig. 4f). Somite desynchronization defects were also observed in a limited number of *Wdr5*^*fl/*+^*;T-Cre* embryos (Fig. 3h). We next intercrossed *Wdr*5^*fl/*+^ and *Rere;T-Cre* mice. Strikingly *Rere*^*om/om*^;*Wdr5*^*fl/*+^*;T-Cre* embryos exhibited a stronger phenotype than either *Rere*^*om/om*^ or *Wdr5*^*fl/*+^*;T-Cre* (Fig. 3l-o) with ~80% (7 out of 9) of the embryos with a right side delay in somitogenesis (Fig. 3k and Supplementary Fig. 4f) and in ~35% (3 out 9) of the embryos, no clear segmented structures forming on the right side (Fig. 3o and Supplementary Fig. 4g,h). This provides strong evidence for genetic interaction between *Wdr5* and *Rere* in the control of somite symmetry. Consistently, transfection of *Rere* and *Wdr5* together in NIH3T3 cells activated RA signalling more strongly than overexpression of either of the proteins alone (Fig. 2l) and *Rere*-dependent RA activation is no longer observed in cells depleted of *Wdr5* (Fig. 2m). Overall, these data reveal the importance of WHHERE in positive regulation of RA signalling and in the control of somitogenesis synchronization in mouse embryos.

Next we analyzed the binding of the WHHERE complex to RA regulated genes by chromatin immunoprecipitation (ChIP). To that end, we first checked the expression kinetics of the RARE-Luciferase reporter and of the endogenous *Rarβ* gene in NIH3T3 cells treated with RA for 1, 2 and 6 hours. At 2 and 6 hours, strong transcriptional activation was observed for both RA targets (Fig. 4a,b). In mouse embryos, we observed by ChIP analysis that the Retinoic Acid Receptor alpha (Rarα) and all the WHHERE complex components were present at the *Rarβ* promoter and at the RARE-containing reporter *RARE-LacZ* (Fig. 4c-f). In ChIP experiments performed 1 hour after RA treatment of NIH3T3 cells, Rere, Wdr5, Hdac1 and Hdac2 but not Rarα increased at the *Rarβ* promoter (Fig. 4g,h). This effect was specific, as no significant enrichment of WHHERE complex members could be observed in regions upstream of the *Rarβ* promoter in similar conditions (Fig. 4h, bottom graph). Recruitment of RNA Polymerase II (Pol II) increased following RA treatment paralleling the WHHERE complex recruitment (Fig. 4i). Then we investigated the requirement of Hdac1 deacetylase activity in transcription activation and Pol II recruitment during RA signalling. Decreased RARE-Luciferase and *Rarβ* expression was observed in TSA-treated cells after RA treatment demonstrating the importance of HDAC activity in early activation of RA target genes (Fig. 4a,b). TSA treatment or siRNA-mediated knockdown of *Hdac1* decreased Pol II recruitment at the *Rarβ* promoter (Fig. 4j,k). No change in Pol II occupancy levels could be detected in RA target genes unresponsive to RA in NIH3T3 cells such as *Cyp26a1* and *Hoxa1* or in promoters of genes unresponsive to RA signalling (Supplementary Fig. 3h-j and data not shown). In transfected cells expressing Flag-Hdac1 alone or Flag-Hdac1 and Rere-HA, we observed binding of Hdac1 to endogenous Rarα. This suggests that Hdac1 could bridge the WHHERE complex to Rarα (Fig. 4l-n). Together, these data support a role of the WHHERE complex in the recruitment of Pol II necessary for early activation of RA regulated genes. This role depends on the deacetylase activity of Hdac1 and Hdac2.

**Figure 4.**
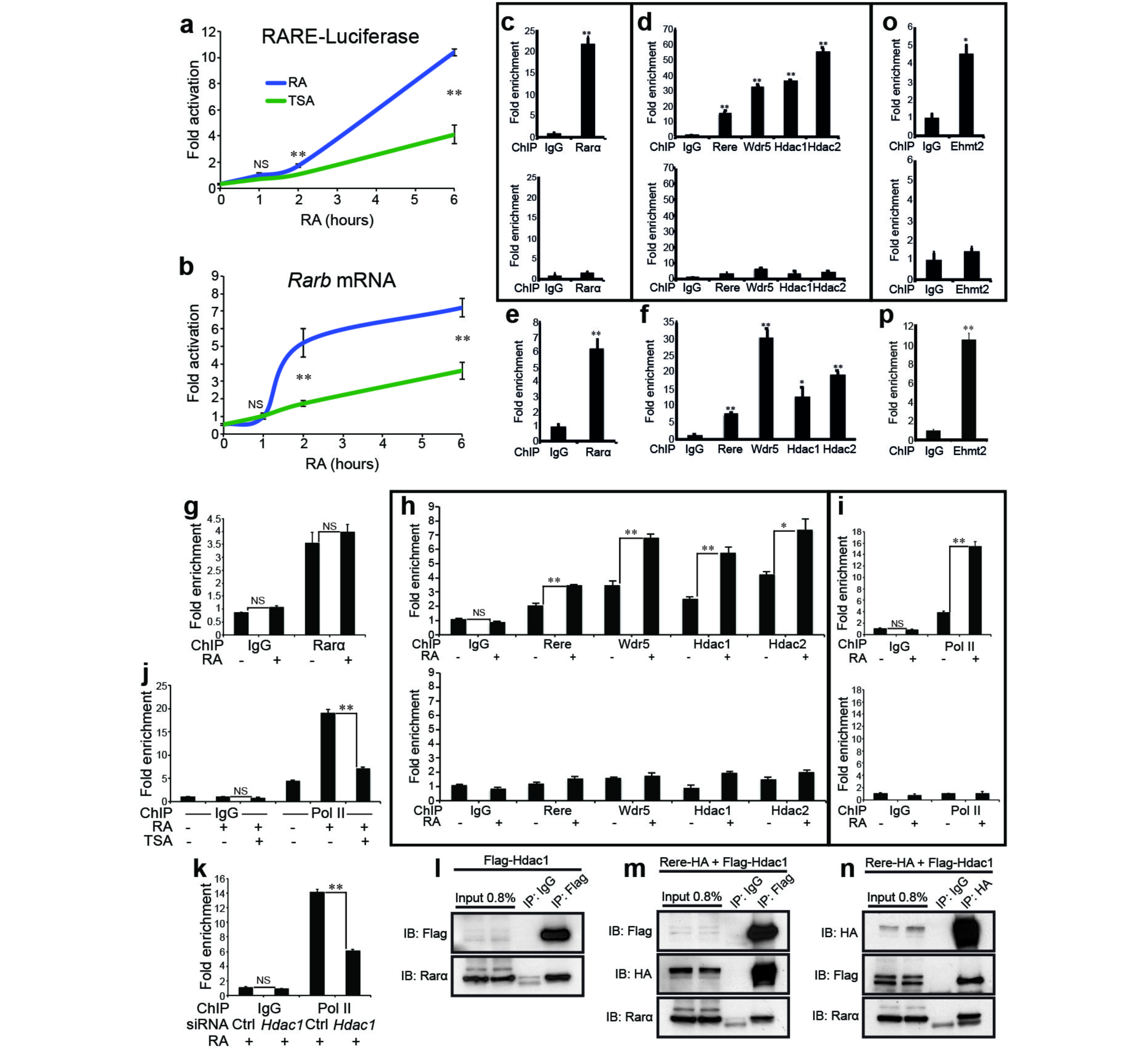
The WHHERE complex is recruited to the promoter of retinoic acid target genes. (**a**) RARE-Luciferase activity of NIH3T3 cells treated with 1 μM RA (blue) or 1 μM RA and 100 nM TSA (green) during 1, 2 or 6 hours (n = 4). (**b**) qPCR analysis *of Rarβ* expression from NIH3T3 cells treated with 1 μM RA (blue) or 1 μM RA and 100 nM TSA (green) during 1, 2 or 6 hours (n = 4). (**c, d**) ChIP analysis of the *Rarβ* promoter from E8.75 RARE-LacZ mouse embryos with specific antibodies against Rarα (**c**) or Rere, Wdr5, Hdac1 and Hdac2 (**d**). Top panels in **c** and **d** show ChIP of the *RARE* element in the *Rarβ* promoter (n = 3). Bottom panels show control ChIP from an upstream region (-3Kb) of the *Rarβ* promoter (data represent mean ± s.d. from triplicate PCR reactions). (**e, f**) ChIP analysis of the *RARE* sequence in the RARE-LacZ reporter with specific antibodies against Rarα (**e**) or Rere, Wdr5, Hdac1 and Hdac2 (**f**) using E8.75 RARE-LacZ mouse embryos (n = 3). (**g-i**) *Rarβ* promoter analysis by ChIP of NIH3T3 cells treated or not during 1 hour with 1 μM RA with specific antibodies for Rar*α* (**g**), Rere, Wdr5, Hdac1 and Hdac2 (**h**) or Pol II (**i**). Top panels in **h** and **i** show ChIP of the *RARE* element in the *Rarβ* promoter (n = 3). Bottom panels show control ChIP from an upstream region (-3Kb) of the *Rarβ* promoter (data represent mean ± s.d. from triplicate PCR reactions). (**j**) ChIP of the *Rarβ* promoter with Pol II antibody using NIH3T3 cells treated with 1 μM RA or 1 μM RA and 100 nM TSA for 1 hour (n = 3). (**k**) ChIP with Pol II antibody from NIH3T3 cells transfected with siRNA for *Hdac1* and treated with 1 μM RA during 1 hour (n = 3). (**l-n**) Co-immunoprecipitation using NIH3T3 cells transfected with expression plasmids for Flag-Hdac1 (**l**) or Rere-HA and Flag-Hdac1 (**m**, **n**). Immunoprecipitation with anti-Flag (**l**, **m**) or anti-HA (**n**) antibody and immunoblots (IB) for Flag-Hdac1, Rere-HA and Rarα. (**o**) ChIP analysis of the *Rarβ* promoter from E8.75 RARE-LacZ mouse embryos with specific antibody against Ehmt2. Top panel in **o** show ChIP of the *RARE* element in the *Rarβ* promoter (n = 3). Bottom panel show control ChIP from an upstream region (-3Kb) of the *Rarβ* promoter (data represent mean ± s.d. from triplicate PCR reactions). (**p**) ChIP analysis of the *RARE* sequence in the RARE-LacZ reporter with specific antibody against Ehmt2 using E8.75 RARE-LacZ mouse embryos (n = 3). In all graphs data represent mean ± s.e.m. unless otherwise specified. NS - not significant, \**P* < 0.05 and \*\**P* < 0.01.

The histone methyltransferase Ehmt2 (G9a) was shown to bind the N-terminal SANT domain of Rere, and together with Hdac1/Hdac2 to regulate the methylation of H3K9 at specific loci leading to the formation of compact heterochromatin and gene silencing^14^. In the proteomic experiment, Ehmt2 (and the related protein Ehmt1) were detected with low NSAF values compared to the members of the WHHERE complex suggesting that its binding to Rere might be transient (Supplementary Fig. 5a). In mouse embryos deficient for *Ehmt2*^23^ crossed to the *RARE-LacZ* reporter^22^, LacZ expression is downregulated suggesting that Ehmt2 is also implicated in positive regulation of RA signalling (Fig. 5a,b). Furthermore, half of the *Ehmt2* mutant embryos presented a desynchronization of somite formation on the right side resembling mutants of members of the WHHERE complex (Fig. 5c-e; Supplementary Fig. 4f). In NIH3T3 cultures, siRNA-mediated knockdown of *Ehmt2* led to a downregulation of the RARE-Luciferase reporter activity (Fig. 5f and Supplementary Fig. 3e), whereas overexpressing Ehmt2 (or Ehmt1) stimulates RA signalling (Fig. 5g and Supplementary Fig. 5b). Co-transfection of *Ehmt2* with *Rere* increases the RA response more than transfection of either one alone (Fig. 5h). siRNA-mediated knockdown of *Ehmt2* inhibited Rere and Hdac1-dependent activation of the RA pathway (Fig. 5i). In NIH3T3 cells, Ehmt2 was recruited at the RARE element of the *Rarβ* promoter after 1 hour of RA treatment while no such enrichment was observed in upstream regions (Fig. 5j and Supplementary Fig. 5c). In mouse embryos Ehmt2 could also be detected at the RARE element present in both the *Rarβ* promoter and the *RARE-LacZ* reporter (Fig. 4o,p). Knockdown of *Ehmt2* in NIH3T3 cells reduced the occupancy levels of Rere, Wdr5, Hdac1, Hdac2 and Pol II at the *Rarβ* promoter (Fig. 5k,l and Supplementary Fig. 5d). In NIH3T3 cells, inhibition of Ehmt2 methyltransferase activity with UNC0638 (U38) or UNC0646 (U46)^24, 25^ did not alter RARE-Luciferase reporter activity or *Rarβ* mRNA expression (Supplementary Fig. 5e,f). Also no difference in H3K9me1 nor H3K9me2 levels was observed by ChIP after 1 hour of RA treatment suggesting that Ehmt2 function is independent from its methyltransferase activity (Supplementary Fig. 5g,h). This suggests that Ehmt2 could function as a scaffold protein to stabilize the WHHERE complex at RA regulated genes to allow Pol II loading. Thus, these results indicate that Ehmt2 acts together with the WHHERE coactivator complex in the RA-dependent control of somite bilateral symmetry.

**Figure 5.**
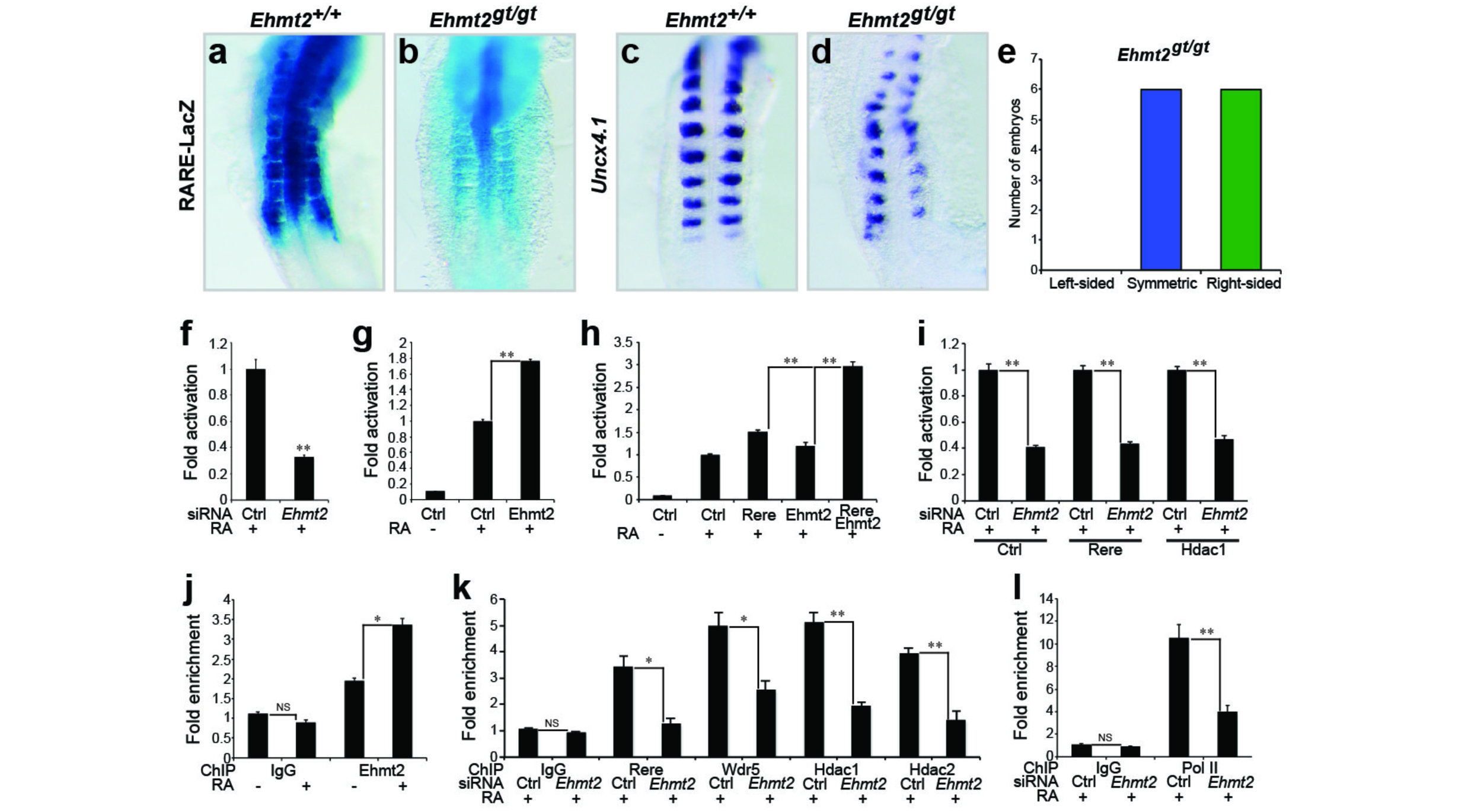
Ehmt2 controls retinoic acid signalling and symmetric somite formation. (**a, b**) RARE-LacZ activity in control *Ehmt2*^+*/*+^ (**a**) and *Ehmt2*^*gt/gt*^ (**b**) embryos at E8.75-E9.0 (dorsal views). (**c, d**) *In situ* hybridization for *Uncx4.1* in wild-type *Ehmt2*^+*/*+^ (**c**) and *Ehmt2*^*gt/gt*^ (**d**) embryos at E8.75-E9.0 (dorsal views). (**e**) Graph representing the number of 7- to 15-somite stage *Ehmt2*^*gt/gt*^ embryos with left-sided(orange), symmetryc (blue) or right-sided (green) delay in somite formation. (**f-i**) RARE-Luciferase activity from NIH3T3 cells treated with or without 1 μM RA for 20 hours. (**f**) Cells treated with siRNA for *Ehmt2* (n = 4). (**g**) Cells transfected with an Ehmt2 expression plasmid (n = 4). (**h**) Cells co-transfected with Rere and Ehmt2 expression plasmids (n = 4). (**i**) Cells overexpressing Rere or Hdac1 and treated with siRNA against *Ehmt2* (n = 4). (**j**) ChIP of the *Rarβ* promoter with a specific antibody for Ehmt2 using NIH3T3 cells treated or not with 1 μM RA during 1 hour (n = 3). (**k, l**) ChIP analysis of the *Rarβ* promoter in NIH3T3 cells transfected with siRNA for *Ehmt2* and treated with 1 μM RA during 1 hour. ChIP was performed with antibodies specific to Rere, Wdr5, Hdac1 and Hdac2 (**k**) or Pol II (**l**) (n = 3). In all graphs data represent mean ± s.e.m‥ NS - not significant, \**P* < 0.05 and \*\**P* < 0.01.

The WHHERE complex contains Wdr5, which is part of complexes such as ATAC, MOF and MLL, that are involved in histone acetylation and H3K4 methylation, which are chromatin modifications associated with transcriptional activation^26^. Despite the requirements of HDAC enzymatic activity for WHHERE function, we observed an increase of the levels of acetylated H3 and H4 as well as H3K27ac on the *Rarβ* promoter upon 1 hour of RA treatment of NIH3T3 cells (Fig. 6a-c). This suggests that Hdac1 and Hdac2 might act on non-histone substrates or both could participate to the stability of the WHHERE complex. The H3K36me3 mark, which is associated to transcription elongation, was also increased (Fig. 6d). We also found that in the same conditions, H3K27me3 is absent from the *Rarβ* promoter (Fig. 6e), whereas H3K4me1 increases while H3K4me2 and H3K4me3 decrease (Fig. 6f-h). MLL3 and MLL4 are complexes which contain Wdr5 and regulate the deposition of the H3K4me1 mark^27^. The MLL3 and MLL4 complexes have been shown to be involved in RA-dependent transcription and Wdr5 might provide a link with the WHHERE complex^28^. Whether these complexes are also required for the WHHERE-dependent Pol II recruitment to the promoter of RA targets remains to be investigated.

**Figure 6.**
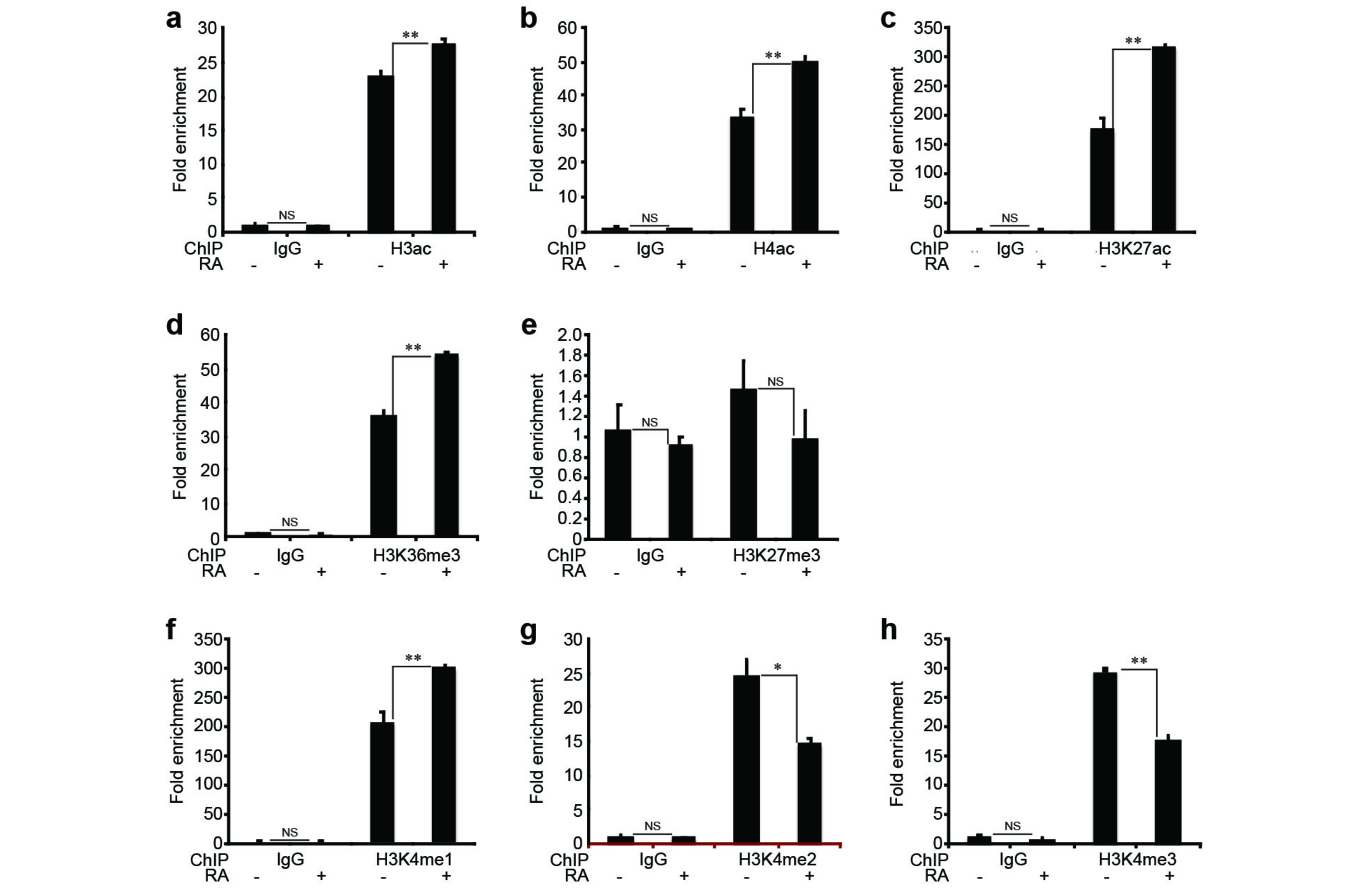
Effect of RA treatment on different histone acetylation and methylation marks. (**a-h**) ChIP analysis of the *Rarβ* promoter from NIH3T3 cells treated with 1 μM RA during 1 hour. ChIP was performed with specific antibodies to H3ac (**a**), H4ac (**b**), H3K27ac (**c**), H3K36me3 (**d**), H3K27me3 (**e**), H3K4me1 (**f**), H3K4me2 (**g**) and H3K4me3 (**h**) (n = 3). In all graphs data represent mean ± s.e.m‥ NS - not significant, \**P* < 0.05 and \*\**P* < 0.01.

The histone acetyltransferase Ep300 (p300) has been shown to acetylate Hdac1 leading to a decrease of its deacetylase activity^29^. In the presence of RA, overexpression of *Ep300* in NIH3T3 cells decreased RA signalling and inhibited Hdac1-dependent RA activation (Fig. 7a). In line with this, transfection of an *Hdac1* mutant form resistant to Ep300 acetylation (H1-6R)^29^ activated more strongly the RA pathway than wild-type Hdac1 (H1-WT) (Fig. 7b). Moreover, Rarα- and Rere-dependent RA activation was inhibited following *Ep300* overexpression whereas transfection of *Kat2a* (*Gcn5*) increased RA signalling more than Rarα or Rere alone (Fig. 7c,d). While treatment of fibroblast cultures with siRNAs against *Rere* or *Kat2a* in the presence of RA decreased RA reporter activity, knockdown of *Ep300* did not affect the RA pathway (Fig.7e). Similarly in *Ep300* mutant embryos (*Ep300*^-*/*-^)^30^, the RARE-LacZ reporter^22^ expression appeared normal (Fig. 7f,g) and somitogenesis progressed symmetrically (Fig. 7h,i). Altogether these results demonstrated that Ep300 negatively regulates Hdac1 activation of RA signalling, and Kat2a can participate together with the WHHERE complex in the activation of the RA pathway.

The molecular mechanisms controlling embryonic bilateral symmetry are still poorly understood. In mouse, the only genes shown to act in this process are *Raldh2* and *Rere* which are involved in the control of RA signalling^7, 8^. Here, we identify a novel protein complex, called WHHERE which associates key epigenetic regulators such as Wdr5, Hdac1 and Hdac2 to the control of bilateral symmetry downstream of RA signalling. In the background of a *situs inversus* mutation (a mutation that can reverse left-right identity of the body), *Raldh2* and *Rere*-deficient mouse embryos can show a reversed somite defect, i.e., left-side delay of somitogenesis^8, 31^. This therefore argues that WHHERE-dependent RA signalling buffers a desynchronizing influence of the left-right determination pathway, involved in asymmetrical development of internal organs. RA signalling was shown to directly antagonize the left identity determinant Fgf8 in the mouse embryo through recruitment of Hdac1 and Prc2, providing a potential explanation for this buffering effect^32, 33^. Thus, our work identifies a new pathway antagonizing FGF signalling downstream of RA to control bilateral symmetry in the mouse embryo.

**Figure 7.**
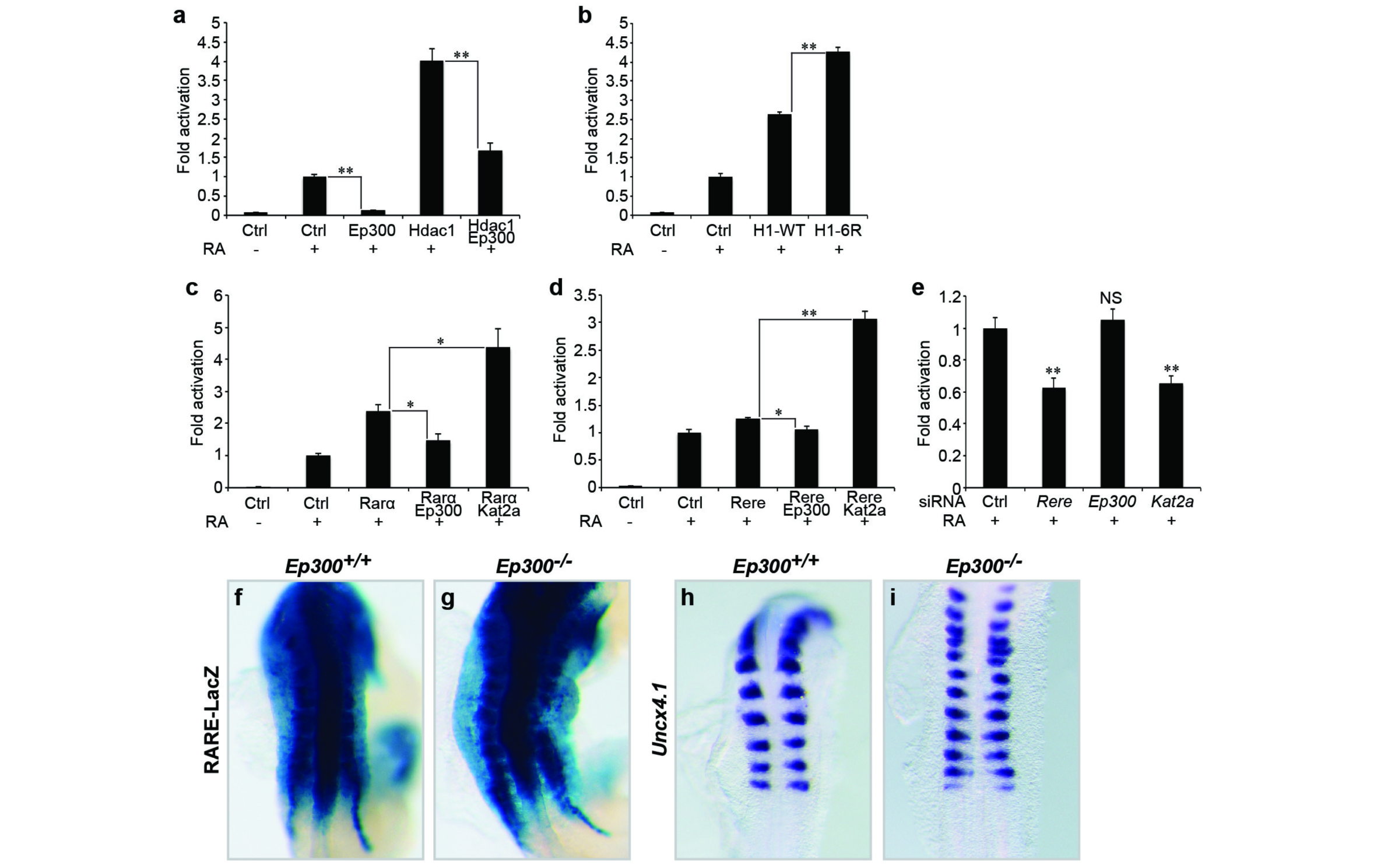
Kat2a but not Ep300 acts as a coactivator for retinoic acid signalling. (**a-e**) RARE-Luciferase activity from NIH3T3 cells treated or not with 1 μM RA for 20 hours. (**a**) Cells transfected with expression plasmids containing *Ep300*, *Hdac1* or both (n = 4). (**b**) Cells transfected with expression plasmids containing human *Hdac1* (H1-WT) or *Hdac1* mutant (H1-6R) (n = 4). (**c**) Cells transfected with expression plasmids containing *Rarα*, *Rarα* and *Ep300* or *Rarα* and *Kat2a* (n = 3). (**d**) Cells transfected with expression plasmids containing *Rere*, *Rere* and *Ep300* or *Rere* and *Kat2a* (n = 3). (**e**) Cells treated either with siRNA for *Rere*, *Ep300* or *Kat2a* (n = 4). In all graphs data represent mean ± s.e.m‥ NS - not significant, \**P* < 0.05 and \*\**P* < 0.01. (**f, g**) RARE-LacZ expression in wild-type *Ep300*^+*/*+^ (**f**) and *Ep300*^-*/*-^ (**g**) embryos at E8.75-E9.0 (dorsal views). (**h, i**) *In situ* hybridization showing somites labeled with *Uncx4.1* in wild-type *Ep300*^+*/*+^ (**h**) and *Ep300*^-*/*-^ (**i**) embryos at E8.75-E9.0 (dorsal views).

Strikingly most of the members of the WHHERE complex including Rere, Hdac1, Hdac2 and also Ehmt2 have been generally associated to transcriptional repression^34-36^. Studies like those reported here may be necessary to fully understand the functions of coregulators, which may act as coactivators or corepressors depending on the developmental context and genes involved. Rere was shown to act as a coactivator for RA signalling^8^ and Hdac1/Hdac2 are required for transcriptional activation of the MMTV promoter downstream of the glucocorticoid receptor^29^. Ehmt2 has also been shown to participate in the positive regulation of a subset of glucocorticoid receptor-regulated genes^37, 38^. Our data shows that in the context of RA signalling these transcriptional repressors can form a complex, which acts as a coactivator of the pathway. The positive role of these proteins in signalling mediated by other nuclear receptors suggests that the WHHERE complex could also have a broader role as a coactivator downstream of nuclear receptor signalling. The positive regulation of RA signalling by Hdac1 and Hdac2 identified in this work might have significant implications for the potential use of HDAC inhibitors for the treatment of RA-sensitive cancers^39^.

## METHODS

Methods and any associated references are available in the online version of the paper.

*Note: Supplementary Information is available in the online version of the paper*.

## ACKNOWLEDGEMENTS

We thank members of the Pourquié laboratory for discussions and comments on the manuscript. We are grateful to R. Schneider, J. Workman, M. E. Torres-Padilla and L. Tora for critical reading of the manuscript. We thank A. Peterson, J. Rossant, E. Olson, R. Feil, M. Lewandoski and Andrew Kung for providing mouse lines. We thank Gordon Hager, Sharon Dent and Richard Eckner for sharing reagents. We thank the Stowers Institute and IGBMC core facilities, particularly the Baculovirus (I. Kolb-Cheynel), Structural Biology (I. Billas) and Cell Culture facilities. We are particularly thankful to M.C. Birling and the Institute Clinique de la Souris for the generation of conditional mouse lines and for animal care (R. Bour and S. Untereiner). The research was supported by the Howard Hughes Medical Institute, the Stowers Institute for Medical Research and the European Research Council (ERC advanced grant).

## AUTHOR CONTRIBUTIONS

G.C.V.-N. designed, performed and analyzed the experiments with O.P‥ G.C.V.-N., M.F. and J.M.G. performed the molecular biology, biochemistry and gel filtration experiments. G.C.V.-N., J.-L.P. and M.M. did the qPCR and qChIP experiments. G.C.V.-N., M.E.S., A.S., L.F. and M.P.W. performed and analyzed the proteomic experiment. G.C.V.-N. did the mouse analysis. G.C.V.-N. and O.P. wrote the manuscript and supervised the project. All authors discussed and agreed on the results and commented on the manuscript.

---

**Supplementary Figure 1.**
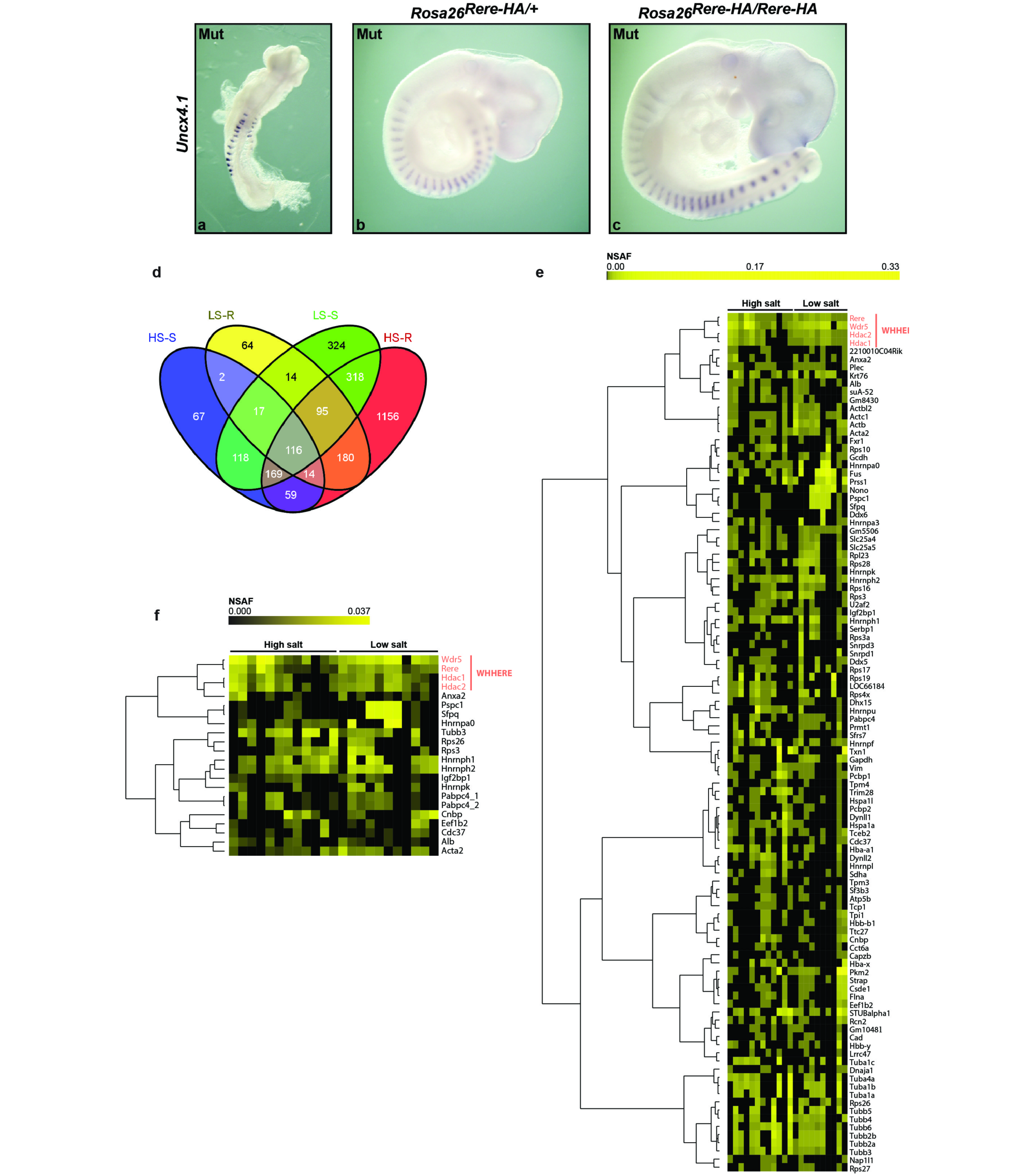
(related to Figure 1) Identification of Rere-associated proteins using multidimensional protein identification technology (MudPIT). (**a-c**) Rescue of *Rere*^*om/om*^ embryos by ubiquitous expression of Rere-HA from the *Rosa26* locus. *In situ* hybridization with *Uncx4.1* in E8.75 *Rere*^*om/om*^ (**a**) (dorsal view), E9.5 *Rere*^*om/om*^*;Rosa26*^*Rere-HA*/+^ (**b**) and E10 *Rere*^*om/om*^*;Rosa26*^*Rere-HA/Rere-HA*^ (**c**) mouse embryos (lateral views). *Rere*^*om/om*^ mutant embryos (Mut). (**d**) Venn diagram comparing the lists of proteins identified after Rere-HA immunoprecipitation in the four different conditions. HS-S: High Salt and Sigma HA beads,LS-R: Low Salt and Roche HA beads, LS-S: Low Salt and Sigma HA beads, HS-R: High Salt and Roche HA beads. The intersection of all four conditions identified 116 distinct entries that correspond to 105 proteins. (**e, f**) Relative protein abundance represented as normalized spectra abundance factor (NSAF) values clustered using Pearson correlation as a distance metric and Ward as a method. Each column represents an individual purification and each row represents a prey protein. The color intensity depicts the protein abundance with the brightest yellow indicating highest abundance and lower intensity indicating decreasing abundance. Black indicates that the protein was not detected in a particular sample. (**e**) Hierarchical clustering of the 105 proteins identified with less stringent criteria. (**f**) Hierarchical clustering of the 22 proteins identified with a more stringent method using the QSpec software. Rere protein clustered with Wdr5, Hdac1 and Hdac2 proteins with the two different methods used for analysis.

**Supplementary Figure 2.**
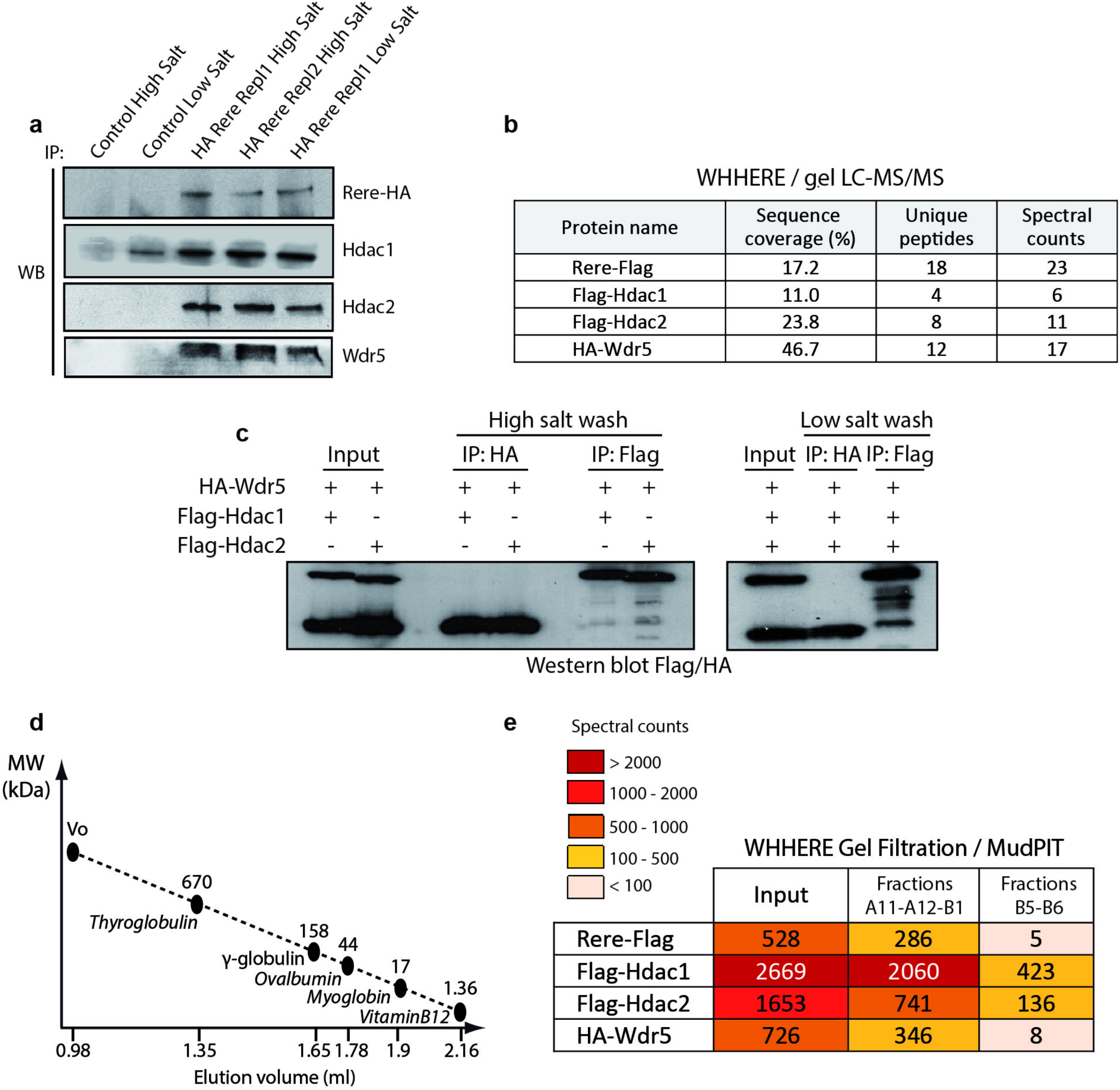
(related to Figure 1) Biochemical characterization of the WHHERE complex. (**a**) Western blot analysis of Rere-HA immunoprecipitation eluates from *T-Cre;RS-Rere-HA* mouse embryos using antibodies against HA, Hdac1, Hdac2 and Wdr5. Replicate 1 (Repl1) and Replicate 2 (Repl2). (**b**) Protein sequence coverage (%), number of unique peptides and spectral counts obtained from the gel LC-MS/MS analysis of Coomassie stained gel bands (shown in Fig. 1c) containing the recombinant WHHERE complex members (Rere-Flag, Flag-Hdac1, Flag-Hdac2 and HA-Wdr5). (**c**) Flag or HA immunoprecipitation from extracts of cells infected with baculoviruses expressing HA-Wdr5, Flag-Hdac1 and Flag-Hdac2 at low (150 mM KCl) and high (500 mM KCl) salt wash. Flag and HA western blot. (**d**) Standard curve performed on the gel filtration column to define the molecular weight (MW) of the WHHERE complex. The MW of Thyroglobulin (670 kDa), y-globulin (158 kDa), Ovalbumin (44 kDa), Myoglobulin (17 kDa) and Vitamin B12 (1355 Da) was plotted based on their elution volume on the Superose 6 column in which the WHHERE complex was analyzed. (**e**) MudPIT analysis of the recombinant WHHERE complex before gel filtration analysis (Input) and after gel filtration from pooled elution fractions containing (A11-A12-B1) or not (B5-B6) the WHHERE complex.

**Supplementary Figure 3.**
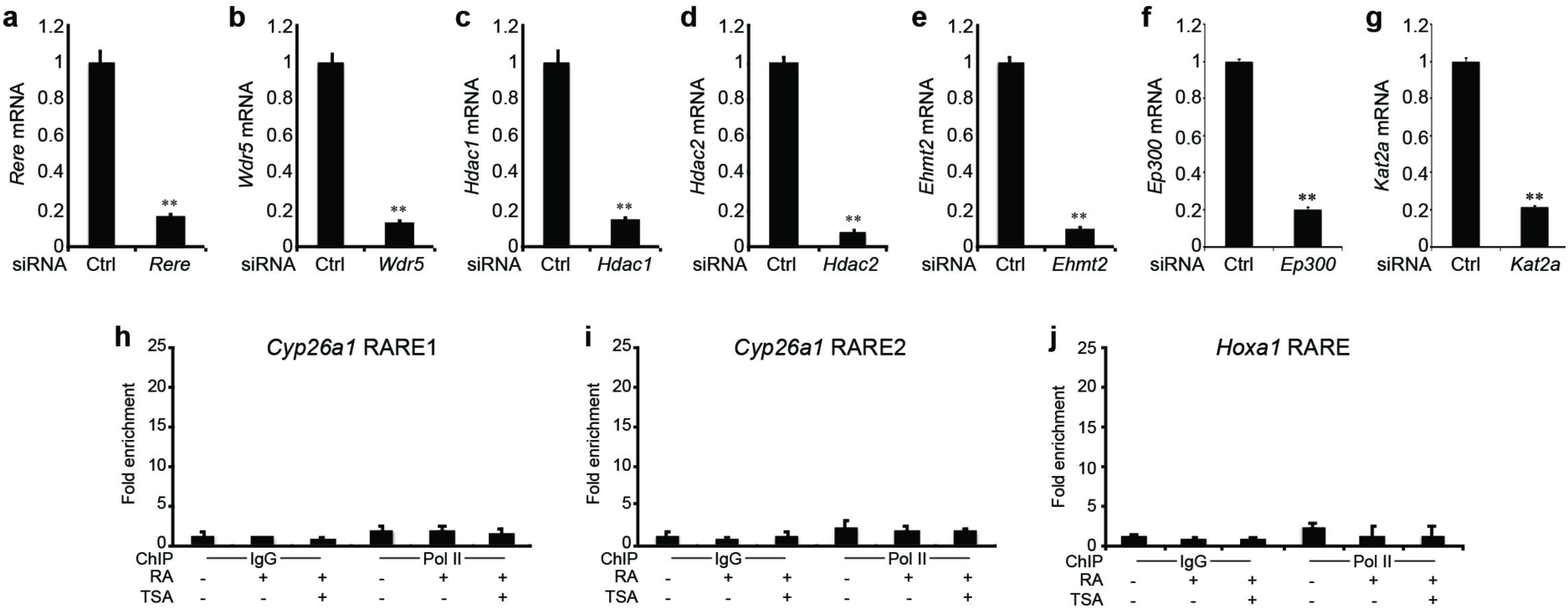
(related to Figures 2, 4, 5 and 7) Efficiency of the siRNAs used to knockdown *Rere*, *Wdr5*, *Hdac1*, *Hdac2*, *Ehmt2*, *Ep300* and *Kat2a*. (**a-e**) Transfection of NIH3T3 cells with siRNAs targeting *Rere* (**a**), *Wdr5* (**b**), *Hdac1* (**c**), *Hdac2* (**d**), *Ehmt2* (**e**), *Ep300* (**f**) and *Kat2a* (**g**). qPCR analysis of *Rere* (**a**), *Wdr5* (**b**), *Hdac1* **c**), *Hdac2* (**d**), *Ehmt2* (**e**), *Ep300* (**f**) and *Kat2a* (**g**) mRNA. Expression level of each gene was normalized to *Rplp0* mRNA (n = 4). (**h-j**) Pol II occupancy on the *Cyp26a1* RARE1, *Cyp26a1* RARE2 and *Hoxa1* RARE elements. ChIP analysis of the *Cyp26a1* RARE1 (**h**), *Cyp26a1* RARE2 (**i**) and *Hoxa1* RARE (**j**) elements with Pol II antibody using NIH3T3 cells treated with 1 μM RA or 1 μM RA and 100 nM TSA for 1 hour (data represent mean ± s.d. from triplicate PCR reactions). In all graphs data represent mean ± s.e.m. unless otherwise specified. \*\**P* < 0.01.

**Supplementary Figure 4.**
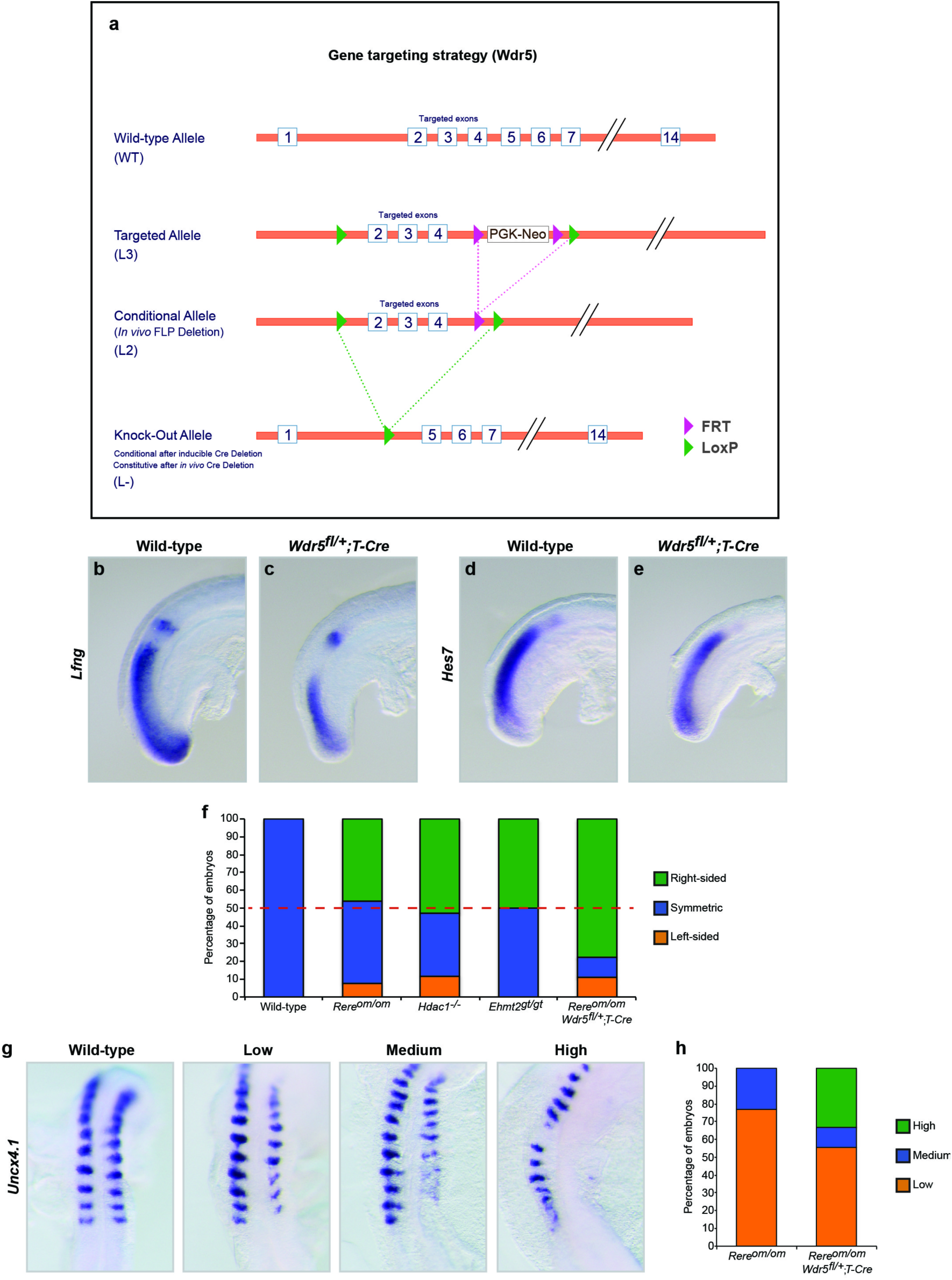
(related to Figures 3 and 5) Asymmetric somite development in mutant embryos for *Rere*, *Hdac1*, *Wdr5* and *Ehmt2*. (**a**) *Wdr5* conditional allele (*Wdr5*^*fl/*+^) mice were generated by inserting two loxP sites flanking the region encompassing exon 2 to exon 4 in C57BL/6N-derived ES cells. The Neo selection cassette flanked by FRT site was removed in vivo using a FlpO deleter mice. The removal of exon 2, 3 and 4 with a Cre deleter mice led to early embryonic lethality before E8.5. (**b-e**) Normal expression of Notch-related genes, *Lfng* and *Hes7*, in *Wdr5*^*fl/*+^*;T-Cre* embryos. *In situ* hybridization for *Lfng* (**b** and **c**) and *Hes7* (**d** and **e**) in wild-type (**b** and **d**) and *Wdr5*^*fl/*+^*;T-Cre* (**c** and **e**) embryos at E9.0-E9.5 (lateral views). (**f**) Graph representing the percentage of 7- to15-somite stage embryos with left-sided (orange), symmetric (blue) or right-sided (green) delay in somite formation in wild-type, *Rere*^*om/om*^, *Hdac1*^-*/*-^, *Ehmt2*^gt/gt^ and *Rere*^*om/om*^*;Wdr5*^*fl/*+^*;T-Cre* embryos. (**g**) *In situ* hybridization for *Uncx4.1* in wild-type and mutant embryos representing different categories of phenotype severity: Low, Medium and High. (**h**) Graph representing the percentage of embryos with Low, Medium and High right-sided phenotype severity in *Rere*^*om/om*^ and *Rere*^*om/om*^; *Wdr5*^*fl/*+^*;T-Cre* embryos.

**Supplementary Figure 5.**
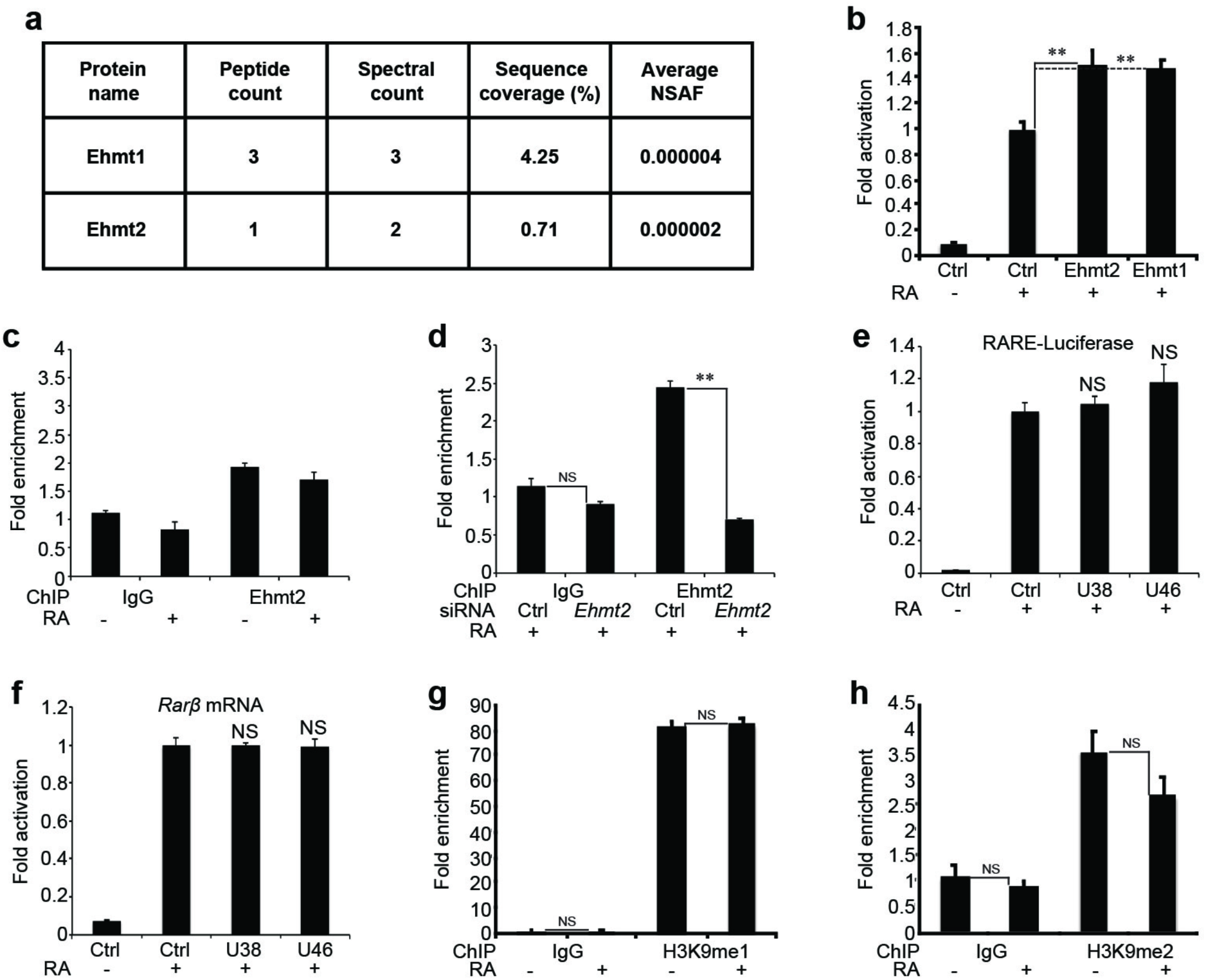
(related to Figure 5) Ehmt2 and the family-related protein Ehmt1 regulate retinoic acid signalling. (**a**) Table representing the peptide count, spectral count, percentage of coverage (%) and average NSAF values for Ehmt1 and Ehmt2 from the MUDPIT analysis of the in vivo Rere-HA immunoprecipitation. (**b**) RARE-Luciferase activity from NIH3T3 cells treated or not with 1 μM RA for 20 hours. Cells were transfected with one of the expression plasmids containing *Ehmt2* or *Ehmt1* (n = 4). (**c**) ChIP analysis of an upstream region (-3 Kb) of the *Rarβ* promoter with a specific antibody for Ehmt2 using NIH3T3 cells treated or not with 1 μM RA during 1 hour (data represent mean ± s.d. from triplicate PCR reactions). (**d**) ChIP analysis of the RARE element in the *Rarβ* promoter using NIH3T3 cells transfected with siRNA for *Ehmt2* and treated with 1 μM RA during 1 hour. ChIP was performed with an antibody specific to Ehmt2 (n = 3). (**e**) RARE-Luciferase activity from NIH3T3 cells treated or not with 1 μM RA for 20 hours and with the Ehmt2 methyltransferase inhibitors UNC0638 (U38) (2μM) or UNC0646 (U46) (2μM) (n = 4). (**f**) *Rarβ* mRNA expression from NIH3T3 cells treated or not with 1 μM RA for 20 hours and with the Ehmt2 methyltransferase inhibitors UNC0638 (U38) (2μM) or UNC0646 (U46) (2μM) (n = 4). (**g, h**) ChIP analysis of the *Rarβ* promoter from NIH3T3 cells treated with 1 μM RA during 1 hour. ChIP was performed with antibodies specific to H3K9me1 (**e**) and H3K9me2 (**f**) (n = 3). In all graphs data represent mean ± s.e.m. unless otherwise specified. NS - not significant, \**P* < 0.05 and \*\**P* < 0.01.

